# SARS-CoV-2 specific plasma cells acquire the phenotype of long-lived plasma cells in the human bone marrow

**DOI:** 10.1101/2022.08.11.503574

**Authors:** Axel R. Schulz, Heike Hirseland, Lisa-Marie Diekmann, Simon Reinke, Sebastian Hardt, Antonia Niedobitek, Henrik E. Mei

## Abstract

Establishment of long-lived plasma cells (PC) in the bone marrow (BM) is important for the development of long-term specific humoral immunity. While SARS-CoV-2-specific, resting, affinity-matured, IgG-secreting plasma cells were described in human bone marrow approx. 6-7 months after infection or vaccination, the long-term durability of these PC remains unclear. We here show that approximately 20% of SARS-CoV-2-specific human BM plasma cells, including RBD-specific PC accommodate the phenotype of long-lived plasma cells, characterized by the lack of CD19 and/or CD45. This result provides evidence in support of the emergence of persistent SARS-CoV-2 specific plasma cells in humans sustaining the durable production of specific serum IgG protecting against severe courses of COVID-19.

---

Long-term serological immunity relies on persistent antigen-specific antibody-secreting plasma cells (PC) and memory B cells, the latter of which can give rise to large numbers of additional plasma cells after re-stimulation by cognate antigen.^1^ While the emergence of specific memory B cells can be analyzed in humans by subsets circulating in the blood, ^2-4^ memory PC do not recirculate, and persist as resting, continuously antibody-secreting cells in the BM for extended periods of time, perhaps for a lifetime. ^1,5,6^

We and others previously characterized human PC lacking the expression of the B cell surface antigen CD19, ^4,7-9^ altogether indicating that CD19^-^ plasma cells are, or comprise, long-lived PC responsible for the observed stability of specific serum antibody titers. ^10^ In support of the exceptional durability of CD19-negative PC, radiocarbon dating of human mucosal lamina propria PC suggested CD19^-^CD45^+^ PC to be approx. 10, CD19^-^CD45^-^ PC 22 years old, while CD19^+^ PC showed contemporary levels of radiocarbon. ^9^ Different from CD19^+^ PC, plasma cells lacking CD19 were enriched in the bone marrow, and mainly secreted IgG relevant for serological immunity, comprised few if any proliferating cells, expressed a pro-survival CD95^-^/Bcl-2^high^ phenotype, and secreted antibodies specific for childhood vaccines. ^4,7^

BMPC specific for SARS-CoV-2, recently described by Turner et al. and Kim et al. ^11,12^ are an important prerequisite for the development of durable SARS-CoV-2 specific humoral immunity, yet their CD19/CD45 status has not been analyzed.

We here assessed the emergence of CD19^-^ and CD45^-^ SARS-CoV-2 specific PC in the human BM in a total of eight femoral head samples (3 men and 5 women, average age 73 years, range 55-86 years) obtained in March-April 2022, by flow cytometry. Seven donors reported having received three or four vaccinations, mostly with mRNA vaccines, against COVID-19. One did not disclose vaccination information. BMPC were detected according to high expression of CD38, and co-expression of antibody kappa or lambda light chains, as described before ^13^ (Figure 1A, S1). High forward scatter signals and expression of CD27 confirmed PC identity (Figure S1). In line with previous studies ^4^ PC made up at average 0.8 % of BM mononuclear cells, and data of approx. 11.000-43.000 plasma cells were available for the analyses from individual donors (Figure 1B, Table S1). S1- and RBD-specific PC were revealed by dual tetramer staining as established before ^14^ and shown in Figures 1A and S2. In line with previous reports employing EliSpot assays, ^11,12^ SARS-CoV-2-specific PC were detectable at frequencies of 0.13% (median, range 0.07-0.67%), comprising at average 64% IgG^+^, 28% IgA^+^ and few if any IgM^+^ BMPC (range 0-2%) (Figure 1C, S3). Antigen-specific PC comprised mainly cells specific for RBD (median 0.09%) next to S1 non-RBD epitopes (0.04%), indicating the selection of potentially neutralizing antibody secreting cells raised against immunodominant SARS-CoV-2 epitopes into the pool of BMPC. Significant frequencies of nucleocapsid-specific PC were detected in only one out of five individuals tested (0.13%, Figure 1C, Table S1) suggesting most individuals were previously vaccinated but not infected with SARS-CoV-2.

**Figure 1.**
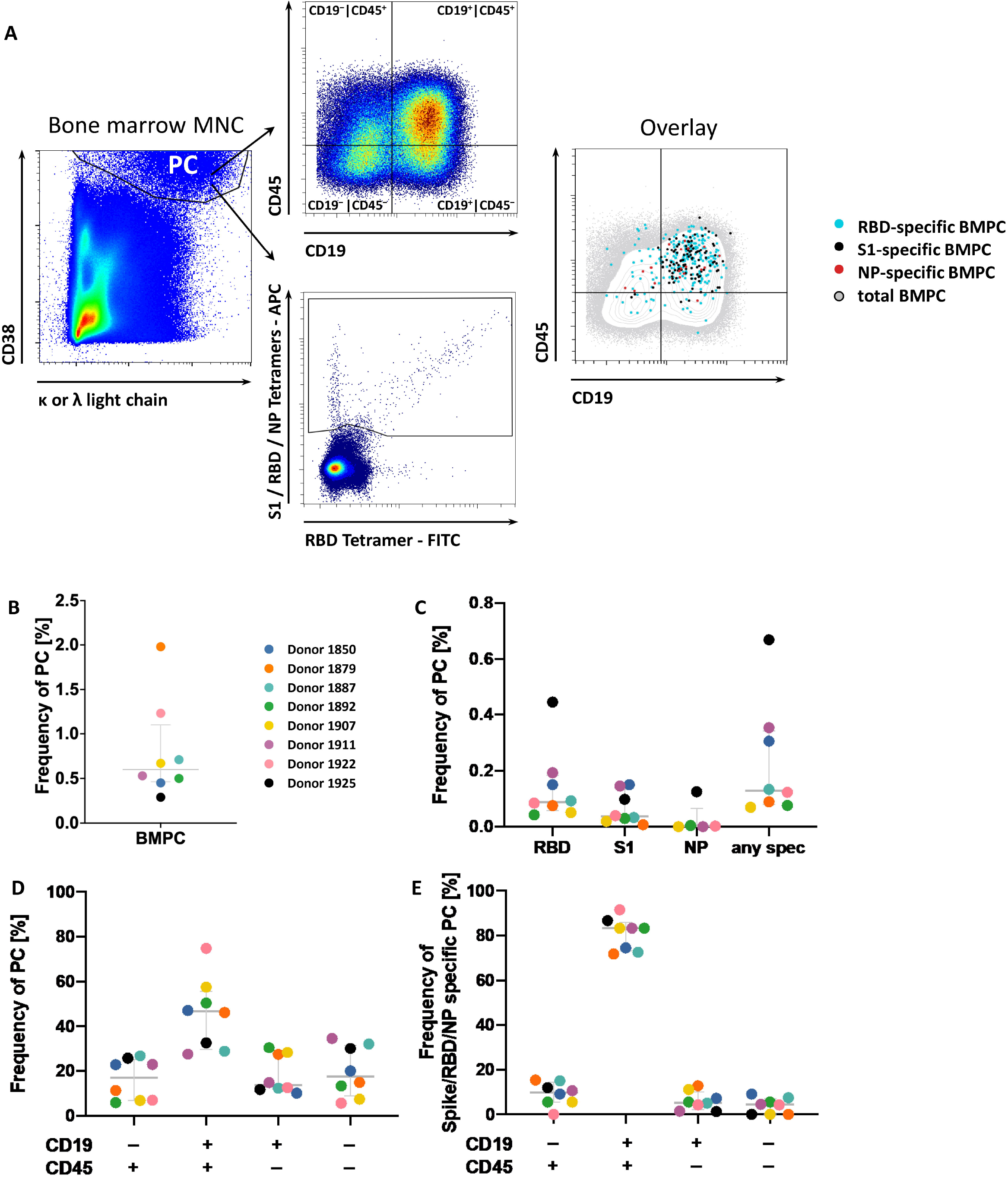
Human bone marrow plasma cell specific of SARS-CoV-2 accommodate the phenotype of long-lived plasma cells. (A) BMPC were analyzed by flow cytometry. PC were gated as shown and analyzed for CD19 and CD45 (top), and antigen specificity (bottom). The overlay comprises concatenated data from eight donors. (B) Frequency of PC in total BM cells. (C) Frequencies of RBD-, S1- and NP-specific and total antigen-specific cells among BMPC. (D,E) PC fractions determined according of CD19/CD45 expression of total and SARS-CoV-2-specific BMPC. Medians and IQR are indicated. PC, plasma cell(s); MNC, mononuclear cells

Total BMPC showed differential expression of CD19 and CD45, confirming previous reports of mature PC, ^4,7,9^ with a median of 47% expressing the conventional CD19^+^CD45^+^ phenotype, next to CD19^-^CD45^+^ (17%), CD19^-^CD45^-^ (18%), and CD19^+^CD45^-^ (14%) PC phenotypes, indicative of advanced maturation and longevity of PC. Significant fractions of SARS-CoV-2 specific PC also expressed these non-canonical PC phenotypes, comprising CD19^-^CD45^+^ (10%), CD19^-^CD45^-^ (4%), and CD19^+^CD45^-^ PC (6%), and thus share the previously described phenotype of long-lived PC (Figure 1D,E).

In paired blood samples available for 5 donors, plasmablasts were detectable at low levels (median frequency, 0.45% of CD19^+^ B cells), comparable to amounts observed during steady-state ^13^ indicating the absence of immune activation at the time of BMPC analysis that may interfere with analyses. A total of four SARS-CoV-2 specific plasmablasts were found in the blood of two out of 5 individuals, while class-switched memory B cells comprised at average of 0.97% SARS-CoV-2 specific cells (Table S1). In addition, only HLA-DR^-^ BMPC were considered in our analysis. Together, this data argues against the possibility that SARS-CoV-2 specific PC observed in the BM samples were from the blood, and support the presence of phenotypically diverse SARS-CoV-2 specific BMPC.

In line with limited antigen experience - 3-4 vaccinations correspond to basic immunization of children according to official recommendations in Germany https://www.rki.de/DE/Content/Infekt/Impfen/Materialien/Downloads-Impfkalender/Impfkalender_Englisch.pdf?blob=publicationFile - SARS-CoV-2 specific PC comprised lower frequencies of the non-canonical, longevity-associated phenotypes compared to overall BMPC (p<0.02, Wilcoxon test). Our data suggest that vaccination against SARS-CoV-2 not only induced SARS-CoV-2-specific BMPC, but that about 20% of these BMPC are destined to last.

It is evident that SARS-CoV-2-specific BMPC of the CD19^+^CD45^+^ phenotype also contribute to serological protection, and their bare presence in the bone marrow indicates that these PC have survived considerable time under steady-state conditions, in the absence of acute plasmablast bursts, as previously discussed. ^4,15^ However, total numbers of CD19^+^ BMPC were shown to be more dynamic, likely reflecting the adaptability of the CD19^+^ plasma cell compartment to emerging challenges. ^4^ Unlike CD19^-^ PC, CD19^+^ BMPC were dependent on the presence of peripheral B cells, ^4^ and only CD19^+^HLA-DR^low^ PC, but not CD19^-^HLA-DR^low^ PC were detectable in the blood after vaccination, presumably as a result of their differential susceptibility to mobilization. ^1,3,4^

This data is in line with the view that humoral SARS-CoV-2 immunity and memory is based on an array of different plasma cell types with different lifespan ^16^ potentially underlying subset-specific regulation. While the regulation of terminal differentiation events leading to the emergence of non-canonical PC phenotypes remains to be revealed, our data show that significant numbers of non-canonical PC can emerge already after timely and quantitatively limited antigen exposure, can be elicited by vaccination, in all likelihood in response to mRNA vaccination, in adults.

Taken together, our results support the early formation of SARS-CoV-2-reactive CD19^+^ and CD19^-^ presumably long-lived plasma cells in the human BM after repeated vaccination against SARS-CoV-2, as an important prerequisite for durable serological immunity preventing severe courses of COVID-19.

## Author contribution statement

ARS, HH, and LMD performed analysis, SR and SH selected donors, provided BM samples and clinical information, AN and ARS established protocols for BM analysis. ARS and HEM made the figures, HEM wrote the manuscript. All authors critically reviewed the manuscript.

## Acknowledgment

The authors wish to thank all participating donors.

## References

1. Yoshida T, Mei H, Dorner T, et al. Memory B and memory plasma cells. Immunol Rev 2010; 237(1): 117–39.

2. Frolich D, Giesecke C, Mei HE, et al. Secondary immunization generates clonally related antigen-specific plasma cells and memory B cells. J Immunol 2010; 185(5): 3103–10.

3. Odendahl M, Mei H, Hoyer BF, et al. Generation of migratory antigen-specific plasma blasts and mobilization of resident plasma cells in a secondary immune response. Blood 2005; 105(4): 1614–21.

4. Mei HE, Wirries I, Frolich D, et al. A unique population of IgG-expressing plasma cells lacking CD19 is enriched in human bone marrow. Blood 2015; 125(11): 1739–48.

5. Manz RA, Radbruch A. Plasma cells for a lifetime? Eur J Immunol 2002; 32(4): 923–7.

6. Slifka MK, Antia R, Whitmire JK, Ahmed R. Humoral immunity due to long-lived plasma cells. Immunity 1998; 8(3): 363–72.

7. Halliley JL, Tipton CM, Liesveld J, et al. Long-Lived Plasma Cells Are Contained within the CD19(-)CD38(hi)CD138(+) Subset in Human Bone Marrow. Immunity 2015; 43(1): 132–45.

8. Bhoj VG, Arhontoulis D, Wertheim G, et al. Persistence of long-lived plasma cells and humoral immunity in individuals responding to CD19-directed CAR T-cell therapy. Blood 2016; 128(3): 360–70.

9. Landsverk OJ, Snir O, Casado RB, et al. Antibody-secreting plasma cells persist for decades in human intestine. J Exp Med 2017; 214(2): 309–17.

10. Amanna IJ, Carlson NE, Slifka MK. Duration of humoral immunity to common viral and vaccine antigens. N Engl J Med 2007; 357(19): 1903–15.

11. Kim W, Zhou JQ, Horvath SC, et al. Germinal centre-driven maturation of B cell response to mRNA vaccination. Nature 2022.

12. Turner JS, Kim W, Kalaidina E, et al. SARS-CoV-2 infection induces long-lived bone marrow plasma cells in humans. Nature 2021; 595(7867): 421–5.

13. Mei HE, Yoshida T, Sime W, et al. Blood-borne human plasma cells in steady state are derived from mucosal immune responses. Blood 2009; 113(11): 2461–9.

14. Cossarizza A, Chang HD, Radbruch A, et al. Guidelines for the use of flow cytometry and cell sorting in immunological studies (third edition). Eur J Immunol 2021; 51(12): 2708–3145.

15. Groves CJ, Carrell J, Grady R, et al. CD19-positive antibody-secreting cells provide immune memory. Blood Adv 2018; 2(22): 3163–76.

16. Amanna IJ, Slifka MK. Mechanisms that determine plasma cell lifespan and the duration of humoral immunity. Immunol Rev 2010; 236: 125–38.

