## Supplemental Figures and Tables for "SARS-CoV-2 specific plasma cells acquire the phenotype of long-lived plasma cells in the human bone marrow"

Supplementary materials

3 figures, 2 tables, experimental details

**Figure S1**

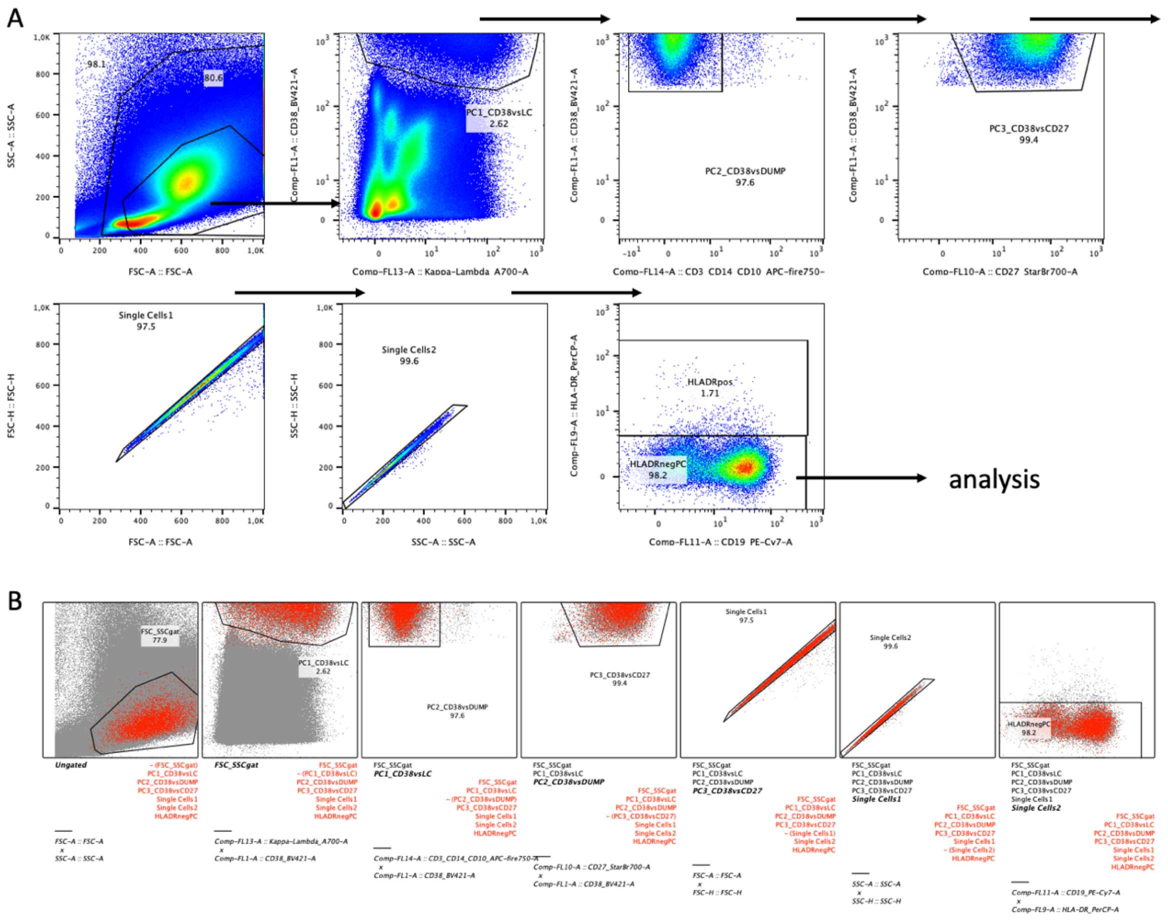


Figure S1. Flow cytometric gating strategy

(A) Total BM cells are shown in the first plot. The large gate was used to determine total numbers of acquired cells, the smaller gate was used to further gate on BMPC according to co-expression of high levels of CD38 and kappa/lambda antibody light chain in the subsequent plot. Additional gates were used to clean up the gated PC by excluding cells stained with antibodies against non-plasma cell lineages (CD3, CD14, CD10), by removing remaining $CD27^{low}CD38^{+}$ cells, cell aggregates in FSC-A vs FSC-H and SSC-A vs SSC-H plots, and finally excluding HLA-DR-expressing cells which may comprise precursors of bona fide PC, that is, plasmablasts (Mei et al., Blood, 2009). (B) Backgating of the same data shown in (A), verifying the above gating strategy. Data of one representative BM sample is shown. Numbers indicate cell frequencies.

**Figure S2**

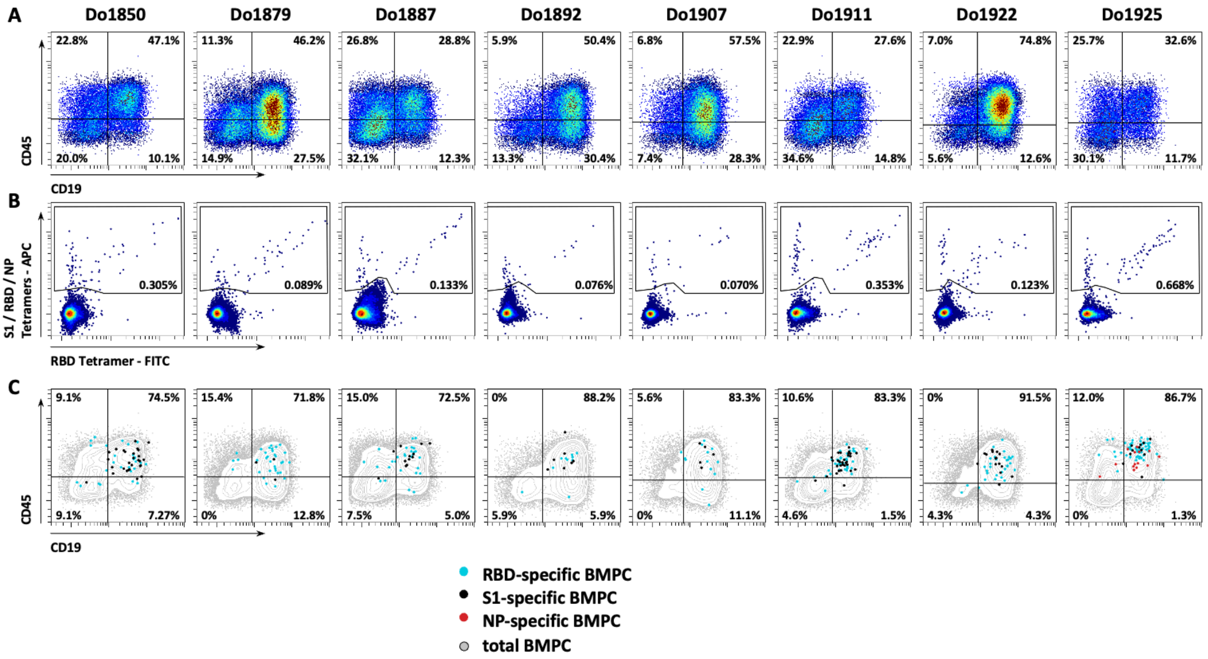


Figure S2. CD19/45 expression by total and antigen-specific BMPC - data of all eight study subjects

BMPC were identified by flow cytometry as shown in Figure 1 and S1. (A) Expression of CD19 and CD45 by BMPC. (B) Detection of SARS-CoV-2-specific PC by dual tetramer staining. (C) Overlay of total and SARS-CoV-2-specific BMPC; the expression of CD19 and CD45 is displayed. Numbers indicate frequencies of gated cells.

Figure S3

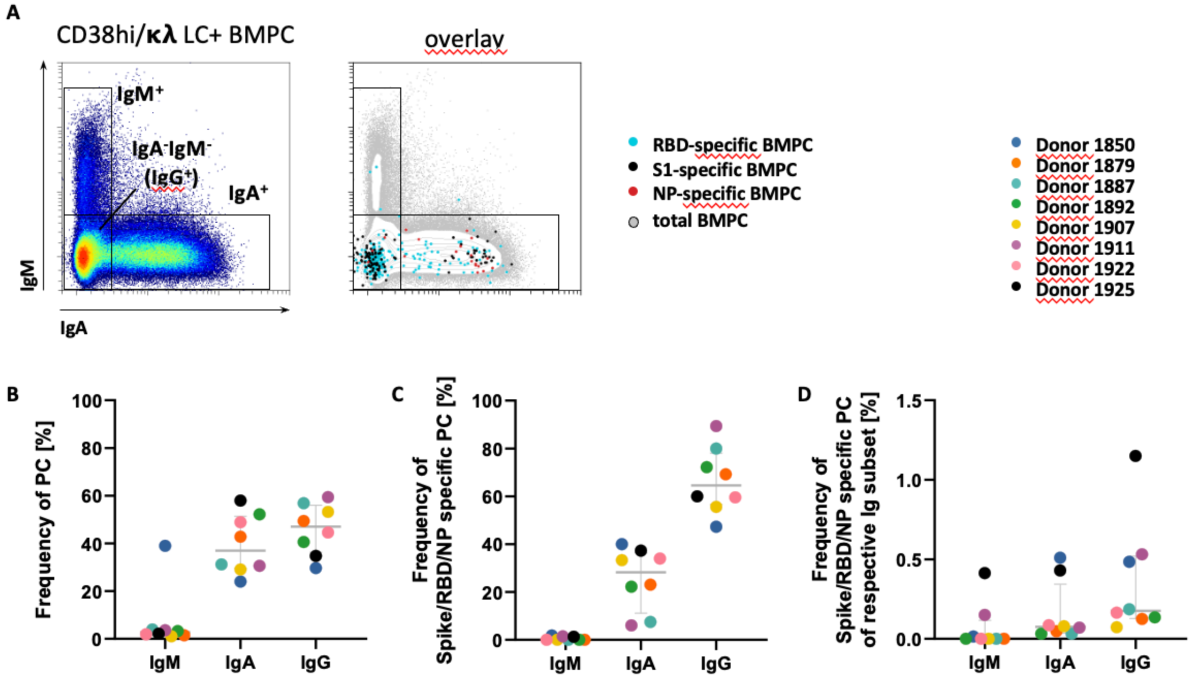


<u>Figure S3. Antibody isotype expression by SARS-CoV-2 specific BMPC</u>

Antibody isotype expression by SARS-CoV-2 specific BMPC was analyzed by flow cytometry. BMPC and SARS-CoV-2 specific BMPC were identified by flow cytometry as shown in Figure 1 and S1, S2. (A) IgM and IgA stainings identify $IgM^+$ and $IgA^+$ BMPC, respectively. $IgA^-IgM^-$ BMPC are considered $IgG^+$ PC, based on previous data (Mei et al., Blood 2015) and scarcity of $IgE^+$ PC in human BM (<1% of PC, unpublished data). The overlay shows concatenated data of all eight study samples. (B) Frequencies of $IgM^+$, $IgA^+$ and $IgG^+$ among total BMPC (C) Frequencies of $IgM^+$, $IgA^+$ and $IgG^+$ among SARS-CoV-2 BMPC (D) Frequencies of SARS-CoV-2 cells among $IgM^+$, $IgA^+$ and $IgG^+$ BMPC. Each dot represents data of one donor. Each donor is colored individually.

### Table S1 Donor and sample characteristics

| BM sample ID | age | sex | self-reported number of vaccinations | self-reported history of COVID-19 infection | use in data | total number of cell events acquired by flow cytometry | total number of plasma cells acquired by flow cytometry | % PC of total cells | specificity stained in BM | antigen-specific PC events | spike antigen (S1+RBD)-specific PC (% of PC) | NP specific BMPC | paired blood sample available | blood PB count in acquired data | specificity stained in Blood | antigen-specific blood PB count in acquired data | % antigen-specific blood PB | % sw-mBC of B cells | blood sw-mBC count in acquired data | antigen-specific blood sw-mBC count in acquired data | % antigen-specific sw-mBC | % plasmablasts among CD19+ B cells |
|---|---|---|---|---|---|---|---|---|---|---|---|---|---|---|---|---|---|---|---|---|---|---|
| 1850 | 80 | female | 3 | no infection | x | $4.01x10^6$ | 18008 | 0.45 | S1+RBD | 55 | 0.31 | n.a. | yes | 378 | S1+RBD | 2 | 0.53 | 10.8 | 8974 | 45 | 0.50 | 0.45 |
| 1879 | 63 | male | not disclosed | not disclosed | x | $2.20x10^6$ | 43637 | 1.98 | S1+RBD | 39 | 0.09 | n.a. | yes | 625 | S1+RBD | 2 | 0.32 | 19.4 | 10748 | 101 | 0.94 | 1.12 |
| 1887 | 80 | female | 4 | no infection | x | $4.26x10^6$ | 30179 | 0.71 | S1+RBD | 40 | 0.13 | n.a. | yes | 144 | S1+RBD | 0 | 0 | 14.8 | 8055 | 136 | 1.69 | 0.26 |
| 1892 | 86 | male | 3 | not disclosed | x | $4.71x10^6$ | 23661 | 0.5 | S1+RBD+NP | 18 | 0.08 | no | yes | 651 | S1+RBD+NF | 0 | 0 | 20.2 | 12721 | 51 | 0.40 | 1.02 |
| 1907 | 67 | male | 3 | not disclosed | x | $3.84x10^6$ | 25831 | 0.67 | S1+RBD+NP | 18 | 0.07 | no | yes | 207 | S1+RBD+NF | 0 | 0 | 12.8 | 8798 | 118 | 1.34 | 0.3 |
| 1911 | 84 | female | 4 | not disclosed | x | $3.51x10^6$ | 18680 | 0.53 | S1+RBD+NP | 66 | 0.35 | no | no | | | | | | | | | |
| 1922 | 55 | female | 3 | not disclosed | x | $3.09x10^6$ | 38065 | 1.23 | S1+RBD+NP | 47 | 0.12 | no | no | | | | | | | | | |
| 1924 | 74 | female | 3 | not disclosed | excluded from analysis, too few total plasma cells | $2.68x10^6$ | 3389 | 0.13 | S1+RBD+NP | 3 | 0.09 | no | no | | | | | | | | | |
| 1925 | 72 | female | 3 | not disclosed | x | $3.86x10^6$ | 11228 | 0.29 | S1+RBD+NP | 75 | 0.67 | yes | no | | | | | | | | | |

n.a. - not applicable
PB - plasmablast
sw-mBC - switched memory B cells, CD19+IgD-IgM- phenotype

Table S2: Flow cytometry reagents

| Target | Fluorochrome | Clone | MQ channel | Dilution | source |
|---|---|---|---|---|---|
| CD38 | BV421 | HIT2 | V1 | 1:100 | Biolegend |
| CD45 | BV510 | HI30 | V2 | 1:200 | Biolegend |
| IgM | BV570 | MHM-88 | V3 | 1:25 | Biolegend |
| CD56 | BV605 | 5.1H11 | V4 | 1:100 | Biolegend |
| Tetramer | BV650 | | V5 | | Biolegend |
| Tetramer | FITC | | B1 | | Biolegend |
| IgA | PE | REA 1014 | B2 | 1:100 | Miltenyi Biotec |
| Tetramer | PEDazzle594 | | B3 | | Biolegend |
| HLA-DR | PerCp | L243 | B4 | 1:100 | Biolegend |
| CD27 | StarBr700 | LT27 | B5 | 1:25 | Bio-Rad |
| CD19 | PECy7 | HIB19 | B6 | 1:100 | Biolegend |
| Tetramer | APC | | R1 | | Biolegend |
| kappa light chain | A700 | MHK-49 | R2 | 1:200 | Biolegend |
| lambda light chain | A700 | MHL-38 | R2 | 1:50 | Biolegend |
| CD3 | APCFire750 | UCHT1 | R3 | 1:100 | Biolegend |
| CD14 | APCFire750 | M5E2 | R3 | 1:100 | Biolegend |
| CD10 | APCFire750 | HI10a | R3 | 1:200 | Biolegend |
| FcBlock | | | | 1:50 | Miltenyi Biotec |
| BV Staining buffer | | | | | Biolegend |

| Tetramer setup | APC | FITC | PeDazzle594 | BV650 |
|---|---|---|---|---|
| S1 | x | | x | |
| RBD | x | x | | |
| NC | x | | | x |

**Supplemental materials and methods**

Samples

Native human bone marrow (BM) was obtained during hip joint replacement surgery, as described before [4]. Information on the donors is summarized in Table S1.

In five cases, paired blood samples were collected prior to surgery in EDTA-coated tubes.

The studies were approved by the ethics committee of the Charité Universitätsmedizin Berlin (application number: EA2/123/21).

Cell isolation from human bone marrow samples

Single-cell suspensions were isolated in a S2 class safety workbench. First, larger fragments of spongeous bone were chopped with a scalpel in sterile petri dish (Corning), if needed. All material was then collected in a 50 mL centrifuge tube (Falcon, BD Biosciences).

PBE (PBS supplemented with 0.2 % BSA and 5mM EDTA) was added to the bone marrow material to liberate cells from bone. After gentle agitation by inverting tube, the suspension was filtered through a 70 µm cell strainer (BD Biosciences), into a new 50 mL centrifuge tube, to remove cell clumps, fat, and solid bone fragments. Larger bone fragments were processed again to recover additional cells.

Afterwards, the cell suspension was centrifuged at 4 °C, 300 × g, for 10 minutes. The supernatant was removed by vacuum. The cell pellet was resuspended in 30 mL PBE.

Mononuclear BM cells were isolated by density gradient centrifugation, by carefully overlaying the filtered BM cell suspension onto 15 mL Ficoll-Paque-Plus (GE healthcare, density, 1.077 g/mL) at room temperature, followed by centrifugation for 20 min at room temperature without brake and low acceleration, at 800 × g. Then, the interphase was collected using a 25 mL serological pipette (Corning) and transferred into a new 50 mL centrifuge tube. After filling up to 50 mL with PBE, the cells were centrifuged at 4 °C, 300 × g, for 10 minutes. The cell pellet was resuspended in 10 mL PBE, and cells were counted volumetrically using a MACSQuant 10 (Miltenyi Biotech). Then, aliquots of $5 \times 10^6$ cells were prepared in 1 mL cryostorage tubes (LVL technologies), pelleted (300 × g, 4 °C, 10 min) and resuspended in 400 µL fetal calf serum (FCS, Corning). Then, 560 µL proteomic stabilizer (Smart Tube Inc, Las Vegas, CA) were added and immediately mixed in by pipetting. After incubation for 12 min at room temperature, tubes were transferred to a -80 °C freezer.

**PBMC isolation from blood**

PBMC were isolated from whole blood by density gradient centrifugation as previously described (Mei et al., Blood, 2015), and further processed as described for bone marrow cell suspensions.

**Flow cytometry**

SARS-CoV-2 spike S1, receptor binding domain (RBD), and for the indicated samples nucleocapsid-streptavidin tetramers (Biolegend and Miltenyi Biotec) were prepared as outlined in Table S2 by incubation for at least 2 h at 4 °C in the dark. Prior mixing with the antibody staining cocktail, tetramer solutions were incubated 1:2 with 40 µM D-Biotin (Thermo Fisher, pre-diluted in PBS) for 5 min at room temperature to saturate unbound streptavidin.

Fixed and cryopreserved PBMC or BM mononuclear cells were thawed for 5 min in cold water (10-15 °C), washed twice in 4 mL PBA buffer (PBS supplemented with 0.5 % BSA and 0.02% sodium azide) pelleted at 700 × g, for 7 min, at 4 °C, resuspended in 2 mL PBA, and counted volumetrically on a MACSQuant 16. Up to $5 \times 10^6$ cells were transferred to a 96 V-bottom plate (Sarstedt), pelleted and resuspended in 15 µL PBA. Next, 25 µL PBA supplemented with 1 µL Fc Blocking solution (Miltenyi) and 0.2 µL heparin (Ratiopharm, Ulm, Germany) were added and cells were incubated for 15 min at room temperature, followed by a 30 min incubation at room temperature in the dark with 25 µL antibody staining cocktail (Table S2). Afterwards, cells were washed twice with up to 200 µL PBA (700 × g, 7 min, 4 °C) and resuspended in 200 µL PBA. 180 µL of the cell suspension were acquired on a MACSQuant 16.

**Data analysis and statistics**

Flow cytometry data were stored in FCS3.1 files and analyzed using FlowJo Version 10 (BD, TreeStar, Ashland, OR) and OMIQ.ai (Santa Clara, CA, USA) by manual gating as shown in Figure S1. Prism version 9 (GraphPad software, San Diego, CA) was used to display data, and perform descriptive statistics and significance testing.
